# Tobacco use and associated factors among Rwandan youth aged 15-34 years: Findings from a nationwide survey, 2013

**DOI:** 10.1101/543843

**Authors:** François Habiyaremye, Samuel Rwunganira, Clarisse Musanabaganwa, Marie Aimée Muhimpundu

**Affiliations:** Department of Institute of HIV/AIDS Diseases Prevention and Control, Non-Communicable Diseases Division, Rwanda Biomedical Center, Kigali, Rwanda; Department of Epidemiology and Biostatistics, College of Medicine and Health Sciences, School of Public Health, Field Epidemiology and Laboratory Training Program, University of Rwanda, Kigali, Rwanda

## Abstract

**Introduction:** Tobacco use is the single most preventable cause of death in the world. The objective of this study was to determine the prevalence of current tobacco use and identify associated factors among Rwandans aged 15-34 years.

**Methods:** This cross-sectional analytical study analysed secondary data collected during the nationally representative Non-Communicable Disease Risk Factors Surveillance survey conducted in 2013 to explore the prevalence of tobacco use in Rwanda and identify factors associated with tobacco use. This study analysed data collected from 3,900 youth participants (15-34 years old), selected using multistage cluster sampling technique. The overall proportion of current smokers, as well as demographic and socioeconomic characteristics of the sample were determined and multivariable logistic regression employed to identify factors independently associated with current tobacco use.

**Results:** The prevalence (weighted) of current tobacco use (all forms) was 8% (95%CI: 7.08-9.01). Prevalence statistically significant was found in the following group: higher prevalence was found among males, young adults aged 24-34, youth with primary school education or less, those from Southern province, people with income (work in public, private organizations and self-employed) and young married adults.

There was no statistically significant difference in prevalence of tobacco use between participants from urban or rural areas (7.8% vs. 8.0%). Factors that were found to be associated with current tobacco use through the multivariate analysis included being a male, aged 25 years and above, having an income, and residing in Eastern, Kigali City and Southern Province compared to Western province.

**Conclusion:** The association between smoking and sociodemographic characteristics among Rwandan youth identified in this study provides an opportunity for policy makers to tailor future policies, and implement coordinated, high-impact interventions to prevent initiation of tobacco use among the youth.

## Introduction

Tobacco use is the single most preventable cause of death in the world[1]. The World Health Organization (WHO) estimates that there are nearly one billion smokers globally[2]. Every year, smoking accounts for more than 7 million preventable deaths worldwide[3]. The annual deaths are expected to reach 8 million by 2030 if no cost effectiveness measures to reduce smoking are initiated [4]. Approximately 80% of all the tobacco attributable deaths occur in low-middle income countries (LMICs)[5] such as Rwanda where tobacco use among adults is estimated to be 13%[6].

Without implementation and enforcement of effective tobacco control policies, smoking prevalence could increase to as high as 22% globally and in the WHO African region by 2030 [7]. The 2013 global burden of disease report estimates that deaths from tobacco use are among the top five causes of mortality in the East African Community (EAC) countries; Burundi, Kenya, Rwanda, South Sudan, Tanzania, and Uganda [8].

Current evidence shows that the path towards smoking and smoking addiction starts at a young age and strongly influences future adult smoking behaviour [9–10]. In Rwanda, no studies have been conducted to identify the major risk factors of tobacco use among the youth. This study was conducted to determine the prevalence of current tobacco use and identify associated factors among Rwandans aged 15-34 years. The Rwandan government policies cap the age of youth as persons aged between 14 to 35 years old[11].

This study was the first comprehensive analysis of the association of current tobacco use and selected socio-demographic characteristics for youth aged 15-34 in Rwanda, and provides evidence for a more targeted programmatic response to tobacco use among the youth in the country.

## Materials and Methods

### Study design and study population

This was a cross sectional analytical study using secondary data collected from the nationally representative Non-Communicable Disease Risk Factors Survey, 2013 of Rwanda.

### Description of the Rwanda Non-Communicable Disease Risk Factors Survey STEPS Survey

The STEPS survey was a population based cross-sectional study conducted in all 30 districts throughout the country from November 2012 to March 2013. The overall objective was to assess the magnitude of risk factors of selected Non-Communicable Diseases in the Rwandan population using the WHO STEPwise approach to surveillance (STEPS).

A multi-stage cluster sampling design was used to select a nationally representative sample. The WHO STEPSwise approach was used to collect data using personal digital assistants (PDAs). These data included socio demographic and behavioural information; physical measurements such as height, weight, blood pressure and waist and hip circumference. Additionally, biochemical measurements were collected to assess total cholesterol, triglycerides levels, fasting blood glucose and urine albumin. In the initial survey, 7200 participants aged 15-64 years were enrolled.

### Data Variables

Information on tobacco use was obtained by asking participants if they were current users of tobacco products. Current smokers were those who had smoked any tobacco product (such as cigarettes, cigars or rolled tobacco) in the previous 12 months. Additional information was collected on behavioral as well as physical and biochemical measurements.

### Statistical analyses

For this secondary study, we extracted data from the STEPS survey for participants aged 15-34 years. Frequencies and percentages were used for descriptive analysis. The primary outcome, current tobacco use among participants aged 15 to 34 years was modeled as a binary variable. We carried out weighted analysis to determine the prevalence of tobacco use and conducted multivariable logistic regression models to identify factors independently associated with current tobacco use. A p-value ≤ 0.05 was considered as significant. We used STATA (StataCorp 11.stata statistical software: Release 12. College Station, Tx:StataCorp LP.) to conduct data analysis.

### Ethical consideration

The survey’s protocol was reviewed and approved by the Rwanda National Ethics Committee (RNEC) and the Centers for Disease Control and Prevention (CDC) Institutional Review Board. Consent was obtained from participants and no individually identifiable information was collected.

## Results

### Characteristics of participants

A total of 3900 participants aged 15-34 years were included in the analysis, of which 2405 (62%) were females. Eighty-three percent (3233) had primary education and below, 80% (944) lived in urban areas, 56% (2187) were married and 80% (3121) were engaged in some form of income generating employment. The sociodemographic characteristics of study participants by tobacco use are shown in table 1.

**Table 1:** Socio-demographic characteristics by tobacco use (N = 3900)

| Variables |  |  | Tobacco use |  |  |
| --- | --- | --- | --- | --- | --- |
| Age group (in years) | N | Percentage | Yes | No | p value |
| 15-24 | 1511 | 38.7 | 4.4 | 95.5 | 0.001*** |
| 25-34 | 2389 | 61.3 | 12.4 | 87.5 |  |
| **Gender** |  |  |  |  |  |
| Men | 1495 | 38.3 | 13.9 | 86 | 0.001*** |
| Women | 2405 | 61.7 | 2.4 | 97.5 |  |
| **Level of Education** |  |  |  |  |  |
| Primary school and below | 3233 | 82.9 | 8.7 | 91.2 | 0.001*** |
| Secondary and High | 669 | 17.1 | 4.7 | 95.2 |  |
| **Province** |  |  |  |  |  |
| Eastern | 972 | 24.9 | 8.8 | 91.1 | 0.001*** |
| Kigali | 560 | 14.3 | 7.2 | 92.8 |  |
| Northern | 651 | 16.6 | 5.4 | 94.5 |  |
| Southern | 773 | 19.8 | 14 | 85.9 |  |
| Western | 944 | 24.2 | 4.3 | 95.6 |  |
| **Employment status** |  |  |  |  |  |
| No earnings | 764 | 20 | 2.7 | 97.2 | 0.001*** |
| Earnings | 3121 | 80 | 9.8 | 90 |  |
| **Residence** |  |  |  |  |  |
| Rural | 2956 | 20 | 8 | 92 | 0.89 NS |
| Urban | 944 | 80 | 7.8 | 92 |  |
| **Marital status** |  |  |  |  |  |
| Married | 2187 | 56.1 | 9.9 | 90 | 0.001*** |
| Others | 1713 | 43.9 | 6.3 | 93.6 |  |

### Tobacco use and associated factors

The prevalence (weighted) of current tobacco use (all forms) was 8% (95%CI: 7.08-9.01). Higher prevalence was found among males, young adults aged 24-34, youth whose highest education was primary school or below, those from Southern province (compared to Western), people with income and young married adults (Table 1). There was no statistically significant difference in prevalence of tobacco use among study participants from urban and those from rural areas (7.8% vs. 8%).

The factors that were found to be associated with current tobacco use after multivariate analysis are shown in table 2. Smoking was associated with being male, aged 25 years and above, residing in Eastern, Kigali City and Southern Province and having an income.

Education attainment was not associated with tobacco use (OR:1.2; 95%CI: [0.8-1.9]).

**Table 2:**
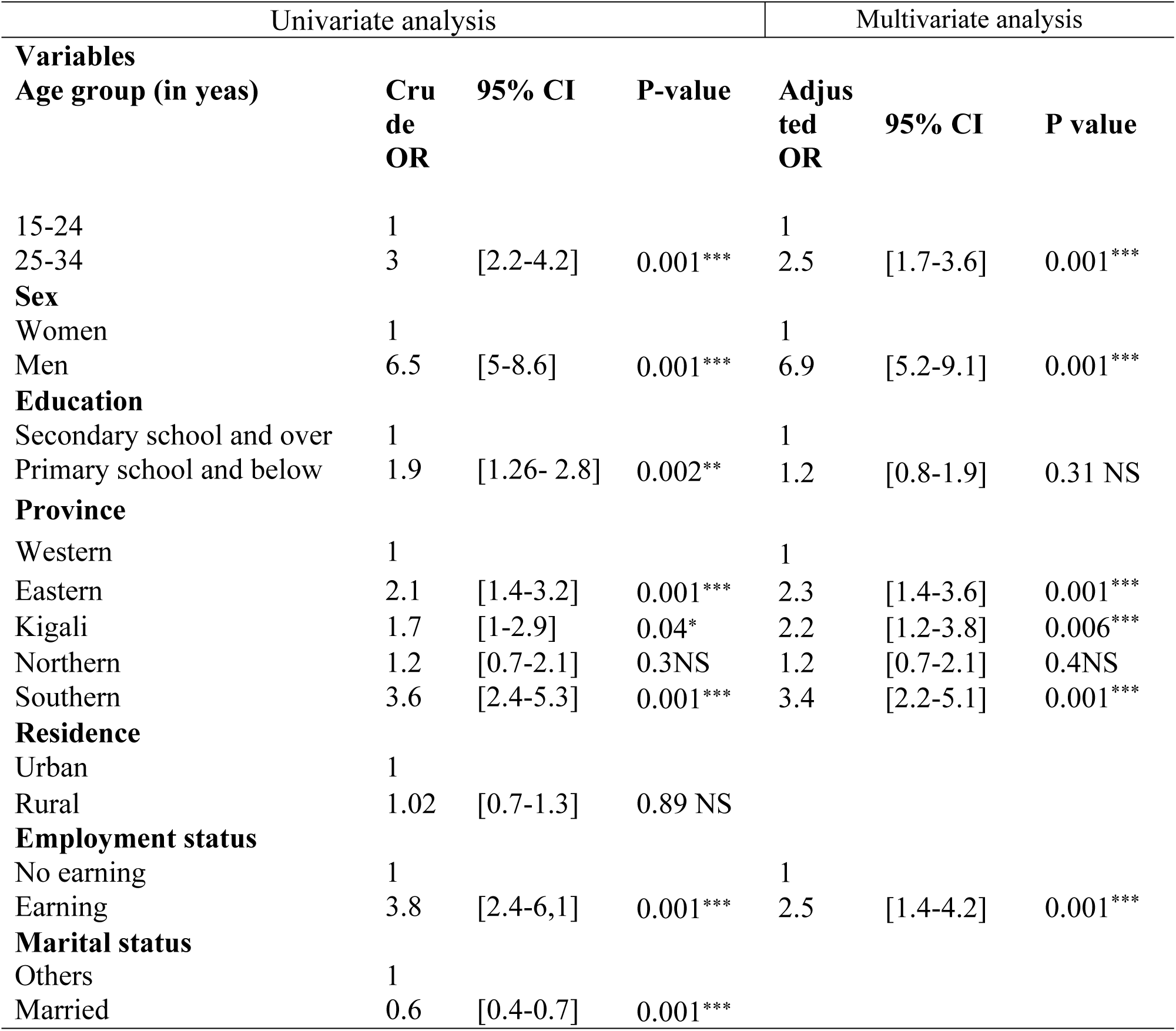
Socio-demographic factors associated with tobacco use among Rwandans aged 15-34 years, 2012-2013.

| Univariate analysis |  |  |  | Multivariate analysis |  |  |
| --- | --- | --- | --- | --- | --- | --- |
| Variables |  |  |  |  |  |  |
| Age group (in yeas) | Cru<br>de<br>OR | 95% CI | P-value | Adjus<br>ted<br>OR | 95% CI | P value |
| 15-24 | 1 |  |  | 1 |  |  |
| 25-34 | 3 | [2.2-4.2] | 0.001*** | 2.5 | [1.7-3.6] | 0.001*** |
| **Sex** |  |  |  |  |  |  |
| Women | 1 |  |  | 1 |  |  |
| Men | 6.5 | [5-8.6] | 0.001*** | 6.9 | [5.2-9.1] | 0.001*** |
| **Education** |  |  |  |  |  |  |
| Secondary school and over | 1 |  |  | 1 |  |  |
| Primary school and below | 1.9 | [1.26- 2.8] | 0.002** | 1.2 | [0.8-1.9] | 0.31 NS |
| **Province** |  |  |  |  |  |  |
| Western | 1 |  |  | 1 |  |  |
| Eastern | 2.1 | [1.4-3.2] | 0.001*** | 2.3 | [1.4-3.6] | 0.001*** |
| Kigali | 1.7 | [1-2.9] | 0.04* | 2.2 | [1.2-3.8] | 0.006*** |
| Northern | 1.2 | [0.7-2.1] | 0.3NS | 1.2 | [0.7-2.1] | 0.4NS |
| Southern | 3.6 | [2.4-5.3] | 0.001*** | 3.4 | [2.2-5.1] | 0.001*** |
| **Residence** |  |  |  |  |  |  |
| Urban | 1 |  |  |  |  |  |
| Rural | 1.02 | [0.7-1.3] | 0.89 NS |  |  |  |
| **Employment status** |  |  |  |  |  |  |
| No earning | 1 |  |  | 1 |  |  |
| Earning | 3.8 | [2.4-6,1] | 0.001*** | 2.5 | [1.4-4.2] | 0.001*** |
| **Marital status** |  |  |  |  |  |  |

|  |  |  |  |
| --- | --- | --- | --- |
| Others | 1 |  |  |
| Married | 0.6 | [0.4-0.7] | 0.001*** |

## Discussion

The WHO STEPS survey was conducted in Rwanda with the aim of providing national level estimates for various NCD risk factors (including tobacco use). To our knowledge, this is the first study conducted to assess tobacco use and associated factors among youth (15-34 years) in Rwanda. Our report using the Rwanda WHO STEPs database provides national-level estimates and information about the prevalence of tobacco use among youth and factors associated with its use in Rwanda. This secondary analysis targeted the youth which comprises over 60% of the Rwanda population and also represents the age at highest risk of tobacco initiation [10–12].

The findings of the study revealed that prevalence of current tobacco use was 2.4% among young women, 14% among young men while the overall prevalence among the youth aged 15-35 was 8%. The results revealed quite similar differences in prevalence by gender among the youth and that of all adults observed during the RDHS 2014/2015 (2% women and 13% men)[14].

These findings are consistent with those of global estimates and other surveys which have found tobacco use to be more prevalent among men than women of all population groups[15]. The observation of higher tobacco consumption among males among Rwandan youth has been consistently observed in many studies conducted in Rwanda and elsewhere. For example, evidence from the Rwanda National health surveys[15][16][13], Rwanda NCDs risk factors surveys[6], the psychoactive substance abuse study[17], Ethiopian study on prevalence of tobacco use and associated factors[18], Southeast Asian Countries study, and the sociodemographic correlates of tobacco consumption in Rural Gujarat, India[19][20], have all found prevalence of tobacco use to be higher among men compared to women. In this case, lower consumption of tobacco use among females in our study, may be accentuated by a social desirability bias.. The culturally tailored stigma associated with tobacco use among females may also have influenced their response.

The lower prevalence of tobacco use among the youth compared to the general population implies that there is a window of opportunity to intervene before the youth begin smoking. Implementing coordinated, high-impact interventions, and stricter implementation of tobacco control measures including mass media campaigns specifically targeting the youth will provide a chance to prevent initiation of tobacco use.

This study identified various socio-demographic factors to be associated with tobacco use. These included age, gender, income status and province of residence. The association with age, income and gender has been observed in multiple studies [16-23]. The association with income was also consistent with findings of the World Health Survey on social determinants of smoking in low and middle-income countries which have shown that smoking is more prevalent among people with income s compared to those without. This has been found to be significant after controlling for age, education and wealth in all settings except women of the low-income country group[24]. In light of existing evidence, persons with higher incomes have the likelihood to avoid smoking initiation and use tobacco less[22]. Nevertheless, considering the present study, I think that income may be a risk factor because tobacco taxation have increased cigarettes prices on the market. Therefore, people in the poorest wealth quintile may not afford tobacco products. These findings contrast with the 2015 study conducted in Ethiopia, which found that adults with low income were more likely to use tobacco as compared to the high income group[18].

This study shows that 8% of Rwandan youth are smokers compared 13% of the general population (15-64 years). This raises concern because young generation will die for tobacco if they don’t quit smoking. This may be attributable to the fact that tobacco industries have end edge technology to market their products and recruit more users among youth. Another aspects is that young people may have social networks/wrong companion to initiate them in tobacco consumption hence they don’t have resources and capacity to avoid initiating tobacco and make necessary steps to quit smoking for those who have already using tobacco. Therefore, behavioral interventions coupled with cessation programs can help these young people to avoid or quit smoking.

This study revealed variation in tobacco use throughout Rwanda’s Provinces. The highest prevalence was found in Eastern Province This difference could be attributed to the availability of contraband cigarettes in this region and tobacco farming at a small scale for consumption purpose.

This study had a number of strengths. First, this was the first nationwide study that allowed the assessing of factors associated with the current tobacco use in Rwandan youth aged 15-34 years. Secondly, the overall response rate of 99.8% for Step 1, and 98.8% for Steps 2 and 3 in the primary study was very high and allowed the findings to be generalizable to all Rwandan youth aged 15-34 years.

The limitations of this study were that although it utilized data from the nationally representative Non-Communicable Disease Risk Factors Surveillance STEPS 2013 of Rwanda, we could not establish a temporal relationship between the associated factors and tobacco use.

In addition, limited variables were collected during the primary data collection and it was not possible to assess other variables for this study. Furthermore, there is possibility of social desirability bias in reporting tobacco use, especially among women and might have led to underestimating of prevalence. Considering that the survey was carried out by health care workers, social desirability might be even higher.

The study has some implications: First, considering the health consequences of tobacco use, having 8% of Rwandan youth as tobacco users represents a substantial risk for morbidity and mortality unless preventive measures are instituted to mitigate the challenge. Cost effective interventions like health education should be prioritized to sensitize the youth on the risks associated with tobacco use. Sustained efforts through price controls and tax measures, comprehensive ban of tobacco smoking in public places, implementation and enforcement of bans to selling tobacco to and by minors in schools and families.

Second, the higher tobacco use among Rwandan youth implies that tobacco initiation occurs at a young age group. Implementing targeted interventions in education institutions (primary and secondary schools, all high learning institutions) should be initiated and strengthened early.

Third, there is a need to establish tobacco cessation program in primary care settings. Tobacco cessation services can be initiated and be made accessible to all who want to quit. Primary health care workers including nurses at health facilities and home based health care workers and teachers can be trained on counselling and other components of tobacco cessation services. Fourth, as tobacco use is a behavioral problem and health workers support is proven to be useful, education of the patients of any non-communicable disease during their contact with health care provider should be a service of immense importance. Additionally, conventional and folk media should be used to educate youth at early ages to prevent initiation of tobacco use.

## Conclusions

The objective of the study was to assess the current smoking prevalence and associated factors among Rwandan youth aged 15-34 years. This study shows that prevalence of tobacco smoking is high among Rwandan youth and is estimated to 8%. The study found that the factors significantly associated with tobacco use among the study population include age, gender, province of residence and employment status.

These findings provide an opportunity for policy makers, decision makers and relevant stakeholders to develop targeted interventions for young people while implementing tobacco control policies and planning tobacco control interventions in general. Since Rwandan youth are at the risk of using tobacco, identifying ways and means of reaching out to these group will be critical to the success or failure of the tobacco control program.

## Acknowledgments

The views expressed in this paper are those of the author(s) and do not necessarily represent the views or policies of the Rwanda Ministry of Health. Authors acknoledge the contributions of Dr. Omolo Jared from the Centers for Disease Control and Prevention (CDC), Kigali, Rwanda; for his inputs and comments to the entire manuscript.

## Author Contributions

Conceived and developed the protocol: FH. Analyzed the data: SR. Wrote the paper: FH. Provided inputs into conception and development of the protocol: FH. Provided inputs and comments into writing of the manuscript: CM, MAM and SR.

